# Temporal topology provides an interpretable framework for neuronal morphogenesis

**DOI:** 10.1101/2025.06.02.657366

**Authors:** Killian Rigaux, André Ferreira Castro, Lida Kanari

## Abstract

Neuronal morphogenesis arises through coordinated neurite dynamics that generate cell-type specific dendritic branching during development. Recent advances in high-throughput time-lapse imaging techniques have transformed our ability to track such growth dynamics, yielding comprehensive anatomical datasets of neuronal morphologies. However, quantifying these structural trajectories during neuronal development remains a major challenge. We introduce the Temporal Topological Morphology Descriptor (TTMD), a framework that combines persistent topology with time-resolved neuronal imaging to quantify developmental trajectories of neuronal architecture. Applying TTMD to datasets of *Drosophila* sensory neurons, we show that developmental stages, branching processes, and mutant-specific growth dynamics can be decoded directly from topology without manual feature selection. Topological morphology descriptor (TMD) accurately resolve major developmental transitions in Class I *Drosophila* neurons, whereas temporal topology (TTMD) is required to uncover subtle alterations in branch dynamics caused by mutations in actin regulatory pathways in Class III *Drosophila* neurons. TTMD further enables automated topological tracking of branch emergence and retraction across development, overcoming the limitations of labor-intensive manual annotation. These results demonstrate that temporal topology provides a unified and interpretable language for neuronal morphogenesis, enabling the automated analysis of developmental neuroanatomy, and identifying pathological alterations in neuronal structure.

## 1 Introduction

The diversity of neuronal cell types reflects the complex developmental processes that shape the nervous system [40, 61]. During development, neurons progress through distinct stages of differentiation, acquiring class-specific morphological features. Dendritic arbors expand and bifurcate to form elaborate tree-like structures that determine synaptic connectivity and influence their biophysical properties [33, 48]. These morphological characteristics are critical for neuronal signal integration and network computation [28, 7]. Understanding how dendrites acquire their cell-type-specific morphological shapes during development is essential to uncovering the relationship between neuronal structure and function, a central goal in systems neuroscience[28, 42].

Although a universally accepted definition of neuronal type remains elusive, neurons are commonly classified based on morphology, spatial location, genetic markers, connectivity, and electrophysiological properties [48, 61, 21]. Among these, morphology serves as an accessible and informative descriptor, reflecting neuronal differentiation and circuit architecture. As such, it offers a valuable entry point for a mechanistic understanding of the development of the nervous system [19, 6, 54, 31, 29, 40, 18, 42]. The emergence of large-scale datasets encompassing diverse dendritic morphologies, enabled by advances in labeling, imaging, and digital reconstruction, has been an essential step in advancing neuroanatomy [50, 5]. However, these expanding datasets have, in turn, raised a fundamental question. How can neuronal morphology be quantified in a robust and efficient manner [49, 23]?

Previous approaches to characterizing neuronal morphology have largely relied on simple morphometric measures, such as neurite length and branching patterns [52, 4]. While informative, these features often capture only narrow, localized aspects of neuronal structure. Moreover, neuroanatomists frequently select morphometrics manually or combine them based on specific research objectives, a process that can introduce bias and overlook a holistic view of neuronal morphology [16]. This limitation becomes increasingly relevant with the advent of time-lapse imaging techniques, which generate high-throughput time series of thousands of labeled neuron images [19, 6, 54, 31, 29]. Because morphological changes during developmental differentiation are often subtle across short time intervals, there is a need for more sensitive and scalable methods capable of detecting fine-grained structural dynamics at high temporal resolution. Therefore, even though conventional techniques have provided valuable information, they cannot fully capture the structural complexity of neuronal development.

Recent advances in the study of neuronal morphologies have enabled the robust classification and clustering of neurons using topological data analysis [35, 34]. The Topological Morphology Descriptor (TMD) represents the branching structure of neuronal trees into a topological descriptor, the persistence barcode [35]. The persistence barcode encodes the start and end radial distances of the neuronal branches from the soma. Therefore, it captures the overall branching structure of neuronal trees into a mathematical descriptor that can be used for detailed analysis and comparison of neurons. The persistence barcode can be alternatively represented as a persistence diagram, a two-dimensional embedding that facilitates averaging across neurons. This representation is useful for generating vectorized versions of the neuronal topology, such as persistence images [1], which can be used as input to classification tasks. These methods have shown promise for static neuronal reconstructions[34, 14]. However, their applicability to time-lapse imaging, where fine-grained morphological changes occur over short intervals, remains an open and largely unexplored question.

Studying the dynamical properties of neurons remains challenging, as it requires time-lapse imaging datasets with high temporal and spatial resolution to track structural changes and link them to neuronal function [30]. The sensory dendritic arborization (da) neurons of *Drosophila* larvae offer a powerful model system to address this challenge, due to their genetic tractability and the ability to perform *in vivo* light-microscopy imaging of entire dendritic arbors through the transparent larval cuticle [19, 6, 54, 33]. Their stereotypical anatomical structure and positional consistency enables their classification into four morphological classes (I–IV) based on increasing dendritic complexity, making them ideal for investigating developmental dynamics and structure-function relationships [43, 47, 53].

To address the limitations of existing morphometric methods in capturing fine-grained structural changes over time, we extended the Topological Morphology Descriptor (TMD) to incorporate temporal dynamics, introducing the Temporal Topological Morphology Descriptor (TTMD). The TTMD of a neuron generates a series of TMD for different time steps. This approach provides a robust quantitative framework for tracking dendritic growth and differentiation over time. We evaluated TMD and TTMD using two time-lapse datasets of Drosophila dendritic arborization (da) sensory neurons. First, we apply TMD to Class I neurons, demonstrating its effectiveness in capturing high-temporal-resolution differentiation processes across embryonic and larval stages. However, TMD alone was insufficient to distinguish more nuanced class-type-specific morphology and altered branching dynamics in Class III neurons from genetically modified mutants. To overcome this limitation, we applied TTMD to Class III neurons to successfully detecte mutant-specific alterations in branching dynamics. Our findings show that the TTMD algorithm provides a scalable, unbiased and computationally efficient alternative to traditional morphometrics, enabling precise identification of developmental stages, mutant phenotypes, and potentially pathological conditions.

## 2 Methods

### 2.1 Time-lapse image series of da neurons morphologies

Time-lapse imaging methods have been described previously in [19, 54], but are summarized here for completeness. Briefly, the flies used in the original studies were kept in standard cornmeal-based food under a 12 hour light / dark cycle at 25^*◦*^*C* and 60% humidity, unless otherwise stated. For time-lapse imaging of dendritic tree structures in dendritic arborization (da) sensory neurons during embryonic and larval stages (L1, L2, and L3), the 221-Gal4 driver was recombined with UAS-mCD8::GFP (Bloomington stock #32187), which labels neuronal membranes with GFP.

In total, 28 *Drosophila* larval da ventral Class I neurons from seven embryos were imaged at 5-minute intervals between 16 hours and approximately 24 hours after egg laying (AEL) (Figure 4). Although this time window spans about 8 hours, due to phototoxicity many neurons died during image acquisition, and some time series were shorter than others. As a result, a continuous 8-hour time series for individual neurons was not technically possible to be acquired [19]. To compensate for these gaps, additional embryos were imaged at different developmental stages to ensure an even distribution of time points across the population. We also note that each embryo contains two ventral Class I da neurons per hemisegment, so multiple neurons were typically imaged per embryo. [22].

Image stacks were acquired at 5-minute intervals; however, due to the high manual burden of dendritic reconstruction (approximately 1–3 days of work per morphology), arbor reconstructions were performed at 30-minute or 1-hour intervals, corresponding to every 6th or 12th image, respectively. All imaging and reconstruction time points are detailed in the accompanying code and metadata (see code availability). From approximately 22.5 hours AEL onward, light peristaltic waves were observed in the embryos, although imaging continued until 24 hours AEL. Following hatching, 20 neurons from five larvae were imaged at discrete developmental time points: 30 hours (first instar), 50 hours (second instar), and 72 hours (third instar) AEL, thereby capturing key stages of larval dendritic development [19].

For each *Drosophila* larva da class III genotype, ten image series were analyzed in wandering third instar larvae. The 30-minute time-lapse sequences were reconstructed every 5 minutes [54].

The confocal image stacks were imported into the TREES Toolbox environment [15], where all dendrites were manually reconstructed using the user interface. In total, the *N*_*classI*_ = 171 and *N*_*classIII*_ = 630 dendrites were reconstructed, defined as the number of neurons imaged multiplied by the number of reconstructed time points per neuron. During the reconstruction process, internode distances (i.e., the spatial resolution of the dendritic structure) were chosen based on total dendritic length: 0.1*µ*m for neurons shorter than 400*µ*m, and 1*µ*m for neurons longer than 400*µ*m.

Time-lapse image stacks were processed as described in the original studies and imported into TREES Toolbox for manual dendritic reconstruction. Each reconstructed dendritic arbor was represented as a tree structure and quality-controlled for common reconstruction errors, including disconnected components and incomplete branching structures using the TREES Toolbox *ver*_*tree*_ function. Each accepted reconstruction was then analyzed with TMD to obtain a persistence diagram encoding dendritic branch topology as a function of radial distance from the soma. Persistence diagrams were vectorized as persistence images, which were used as input features for downstream visualization and classification analyses.

The term “developmental stages” refers to biologically defined phases of neuronal maturation rather than arbitrary positions along a temporal axis. In Drosophila, embryonic and larval stages represent tightly regulated developmental transitions controlled by genetic programs and accompanied by substantial changes in neuronal morphology and dendritic arbor organization [6, 19]. The Class I dataset follows neurons across embryonic and larval development, enabling us to investigate whether our analysis captures these genetically regulated phases of morphological differentiation.

Class III dataset addresses a distinct biological question. Neurons were examined within a single developmental stage but across different genetic backgrounds that affect actin-dependent dendritic branch dynamics. Consequently, the 5-minute intervals in the Class III time-lapse sequences do not represent transitions between developmental stages, but instead capture short-timescale branch dynamics within a fixed developmental context. In this analysis, genotype or mutant condition serves as the biological label, while the temporal ordering of the sequences is incorporated into the topological feature representation to characterize dynamic differences between genotypes.

### 2.2 Preprocessing of data

We first tested the quality of dendritic reconstructions for common errors, such as disconnected components and completeness of branching processes. 22 reconstructed class I dendrites exhibited a very low number of branch points, which indicated that the corresponding dendrite morphologies had not differentiated yet. The presence of such a feature therefore, prevents the use of the TMD to classify them, due to possible incompleteness of the dendritic processes.

Thus, a reduced dataset that excludes the 22 non-differentiated class I neurons (that concern *t5_30, t6_30*, and *tl_nov_friday* series) was considered for the subsequent analysis. It comprises *N*_*classI*_ = 149 and *N*_*classIII*_ = 630 dendrites. The elements of this set are also referred to as “neurons”.

### 2.3 Topological Morphology Descriptor

Persistent homology [62] captures the topological properties of shapes across different scales. It has been used successfully for a variety of tasks including the reconstruction, recognition, and matching of objects. Persistence homology captures relevant information on the underlying shapes by pairing critical values of a function, represented as points in a 2D diagram, the persistence diagram. Typically the function used as a filtration in persistence homology is the height function [62, 9]. In [35] they introduced an alternative algorithm to compute persistence homology by generating a topological descriptor of neurons (Topological Morphology Descriptor, TMD) based on radial distances from the soma, which produces a persistence barcode from any tree-like structure.

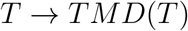

The topological morphology descriptor (TMD) [35, 34] transforms tree-like structures, such as neuronal morphologies, with a real-valued function *f* : *V* → ℝ on the tree nodes, to a persistence barcode through a filtration function. It produces an embedding of the graph in a two-dimensional space that has been shown to capture well the topological properties of the graph and has been used for the classification and clustering of rodent [34] and human [17, 37] pyramidal cells. Each branch within the tree is represented by a line, defined as the topological lifetime of a branch, in the barcode which encodes the first and the last “time” (in units of the function *f*) that the branch was detected in the tree structure. Note, that the start and end of a branch here refers to a single time point, and not the beginning and end positions of the same branch across time, during the time-lapse sequence. The persistence barcode is a collection of lines that encode the topological lifetime of each branch in the underlying tree structure. Alternatively, the start and end “times” of the branches can be represented as two-dimensional points in a persistence diagram. The two representations are equivalent and will be used interchangeably in the manuscript. An example embedding is presented in Figure Figure 1.

**Figure 1:**
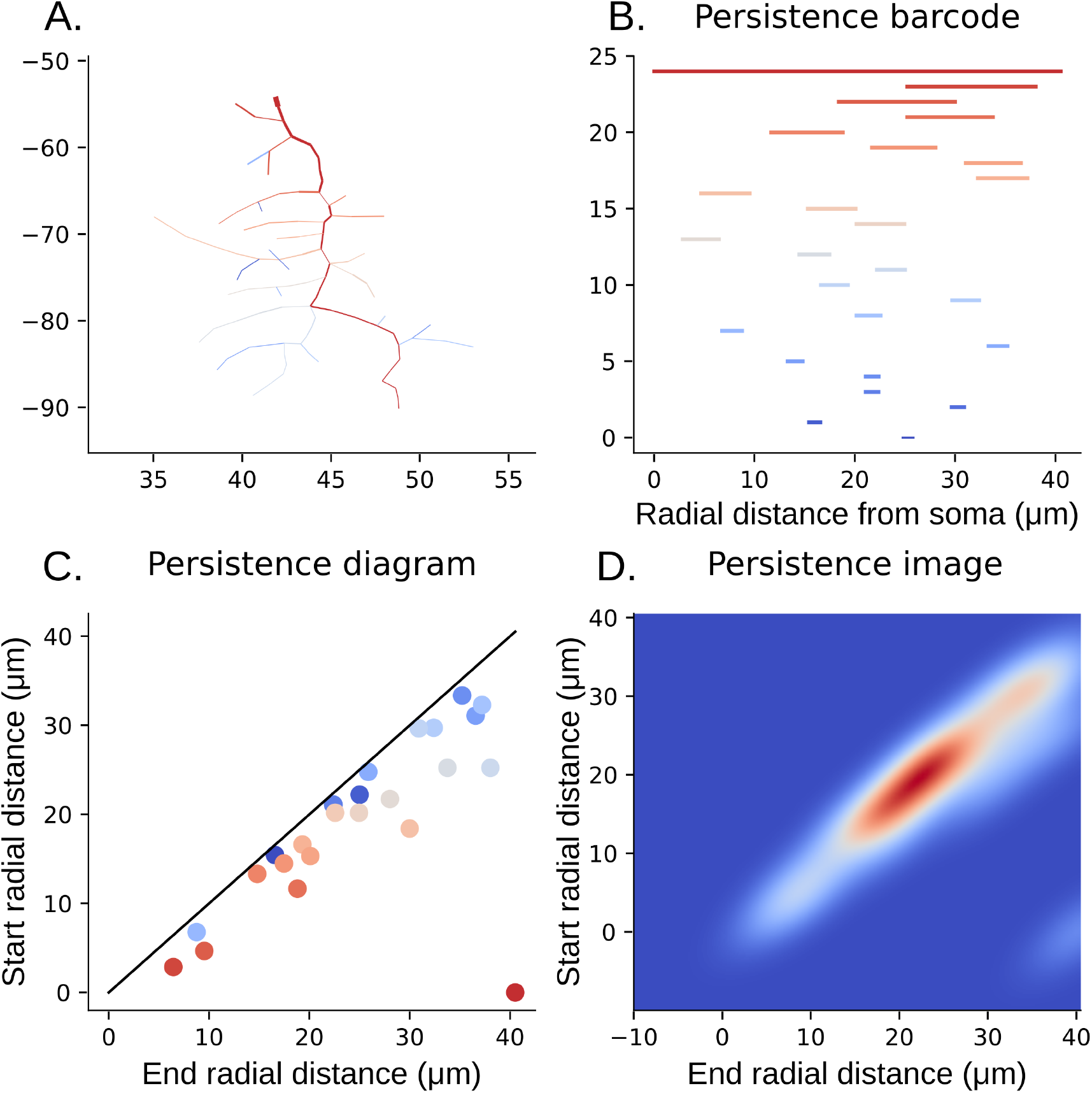
Topological Morphology Descriptor extracted from a Class I neuron. **A**. The reconstructed c1da dendritic tree. Colors represent the branches(tree components from a bifurcation to the terminal tip)and the bars in the persistence barcode, ranging from longer (red) to smaller branches (blue). **B**. The TMD barcode is produced through filtration of the apical graph using radial distances. **C**. The persistence diagram presents the same information plotted in two-dimensional space, where axes represent the birth and the death of each segment of the bar code. **D**. The persistence image shows a Gaussian kernel density estimate applied to the persistence diagram in C.

### 2.4 Classification process

For the classification process, we extracted the TMD of all neurons and vectorized them as persistence images. We associated each persistence image with expert-defined labels. Supervised learning is then performed on this data using the *leave_one_out_mixing* method. We selected the classifiers *LinearDiscriminantAnalysis, QuadraticDiscriminant Analysis* and *DecisionTreeClassifier* of the *scikit-learn* Python library. The linear classifier separates data with a linear boundary, whereas the non-linear boundary of the quadratic one offers a finer distinction. The decision tree classifier assigns data to classes based on binary decisions, and is the most advanced classifier that we used. We decided to keep the classifier selection simple to optimize training times.

The *leave_one_out_mixing* method evaluates classification quality using leave-one-out validation in order to reduce potential overfitting effects. In this approach, the dataset is divided into n subsets corresponding to individual samples. For each iteration, the classifier was trained on *n* − 1 samples and tested on the remaining unseen sample. This procedure was repeated *n* times so that each sample is used once as test data. We adopted this strategy instead of a conventional k-fold cross-validation because the relatively small sample size would produce training and testing subsets that are too small for stable classifier performance.

The *leave_one_out_mixing* results are summarized through a confusion matrix, which reports the classification success rate for each class. Overall classification performance is quantified using the accuracy, computed as the ratio of correct predictions to the total number of predictions. We do not report balanced accuracy because the number of samples across classes is relatively balanced.

### 2.5 TTMD Classification process

To incorporate temporal structure into the classification framework, we developed TTMD, which generates one persistence diagram per neuron at each of the *t* recorded time points. This yields a longitudinal topological representation of neuronal morphology across development. We evaluated the impact of temporal information using three classification strategies. All classification with TTMD was performed using a decision tree classifier.

#### 1. Independent-timepoint classification (“no memory”)

At each time point *t*_*k*_, we trained and evaluated a classifier using only the persistence diagrams computed at *t*_*k*_. Each time point was treated as an independent dataset. This approach isolates morphology-specific discriminative structure at distinct developmental stages and allows identification of stages at which neuronal classes are most separable.

#### 2. Cumulative classification (“with memory”)

At time point *t*_*k*_, the training set included persistence diagrams from all time points {*t*_1_, …, *t*_*k*_*}*. Diagrams corresponding to the same neuron across time shared the same class label. This cumulative strategy tests whether progressive incorporation of earlier developmental information improves class separability and quantifies how many developmental stages are required before reliable discrimination emerges.

#### 3. Full-dataset classification (“all data”)

In this condition, classifiers were trained using persistence diagrams from all neurons across all time points simultaneously. This represents the upper bound of performance achievable when complete temporal information is available.

Together, these three strategies disentangle stage-specific discriminability from temporally integrated discriminability and allow assessment of whether developmental trajectory information provides additive classification value beyond single-timepoint morphology.

### 2.6 Temporal Topological Morphology Descriptor

Some datasets depend on several parameters, and in this case, we are interested in studying the persistent homology of a multiparameter family of spaces [25]. Persistent homology for one-parameter datasets is well studied, but this is not the case for multiparameter persistence homology, which is often hard to handle. For the topological analysis of systems that vary across multiple dimensions, several methods have been proposed. One prominent approach is multipersistence [10, 51, 25], which extends persistent homology to incorporate multiple parameters. Multipersistence has been applied in diverse fields, including the analysis of time series [38] and mitochondrial image analysis [12].

In the context of growing neurons, multipersistence could capture variations across spatial dimensions while integrating temporal information. However, this method is computationally expensive and relies on continuous temporal data, which is not available for biological neurons due to the challenges of continuous tracking and their inherent complexity. Consequently, an alternative approach is required to effectively describe the growth dynamics of neurons.

Therefore, we define the Temporal Topological Morphology Descriptor, *TTMD*, as the collection of TMD representations of a neuronal tree at different time steps

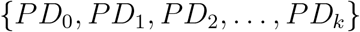

. At each time step, the TMD algorithm is applied independently to the reconstructed morphology of the respective time step.

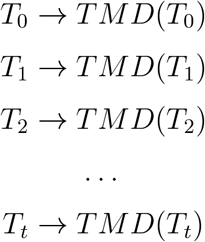

However, the TTMD does not yet establish a clear connection between timepoints.

For the classification tasks in this study, we used the set of persistence images from all time points as a combined feature vector, see also Section 2.4.

### 2.7 TMD - vineyards

Another approach to study datasets that vary across a dimension is the persistence homology transform [57] and the persistence vineyards [13, 27, 56]. Typically used to track changes in persistence across a varying parameter, such as different viewing angles of an object [57, 56], persistence vineyards have also been proposed to study the development of neuronal morphologies [41]. We extend the vineyard framework to analyze developmental differences in growing Drosophila neurons. Specifically, we modify the traditional definition of vineyards to establish correspondences between persistence diagrams extracted at different time points using the Topological Morphology Descriptor (TMD) [35, 34].

In our use case of developing neuronal trees, we initially treat the trees at different time steps as independent and extract their persistence diagrams according to the TMD algorithm [35]. Given a neuron *n* at different times *n*_1_, *n*_2_, …, *n*_*t*_, we represent each tree by a respective diagram:

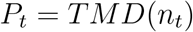

#### 2.7.1 Matching algorithm

To compare two consecutive time steps *t*_1_, *t*_2_, the points in the respective persistence diagrams *P*_1_ and *P*_2_ can be matched to each other. The points in the respective diagrams *P*_1_, *P*_2_ are matched using the Munkres algorithm [46] (also known as the Hungarian or Kuhn-Munkres algorithm) to solve the optimal assignment problem between diagram points. Given two persistence diagrams *P*_1_ and *P*_2_, each represented as a set of birth-death pairs, the algorithm computes a bijection

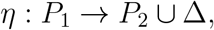

where Δ denotes the diagonal, allowing unmatched features to be paired with the diagonal.

A cost matrix is constructed in which each entry corresponds to the distance (e.g. *L*_∞_ or *L*_*p*_) between a point in *P*_1_ and a point in *P*_2_, or to the cost of matching a point to the diagonal, which represents the appearance or disappearance of a topological feature. The Munkres algorithm then finds the assignment that minimizes the total matching cost, yielding the optimal correspondence between topological features. This procedure runs in 𝒪 (*n*^3^) time, where *n* is the total number of points in the two diagrams, and forms the basis for computing bottleneck or Wasserstein distances as well as for tracking feature evolution in persistence vineyards.

By matching each point in the persistence diagram of a time step *t*_1_ with the next time step *t*_1_ we generate a sequence of points across time to track the evolution of each branch in the neuronal tree (Figure 2), creating piecewise linear curves across the time steps.

**Figure 2:**
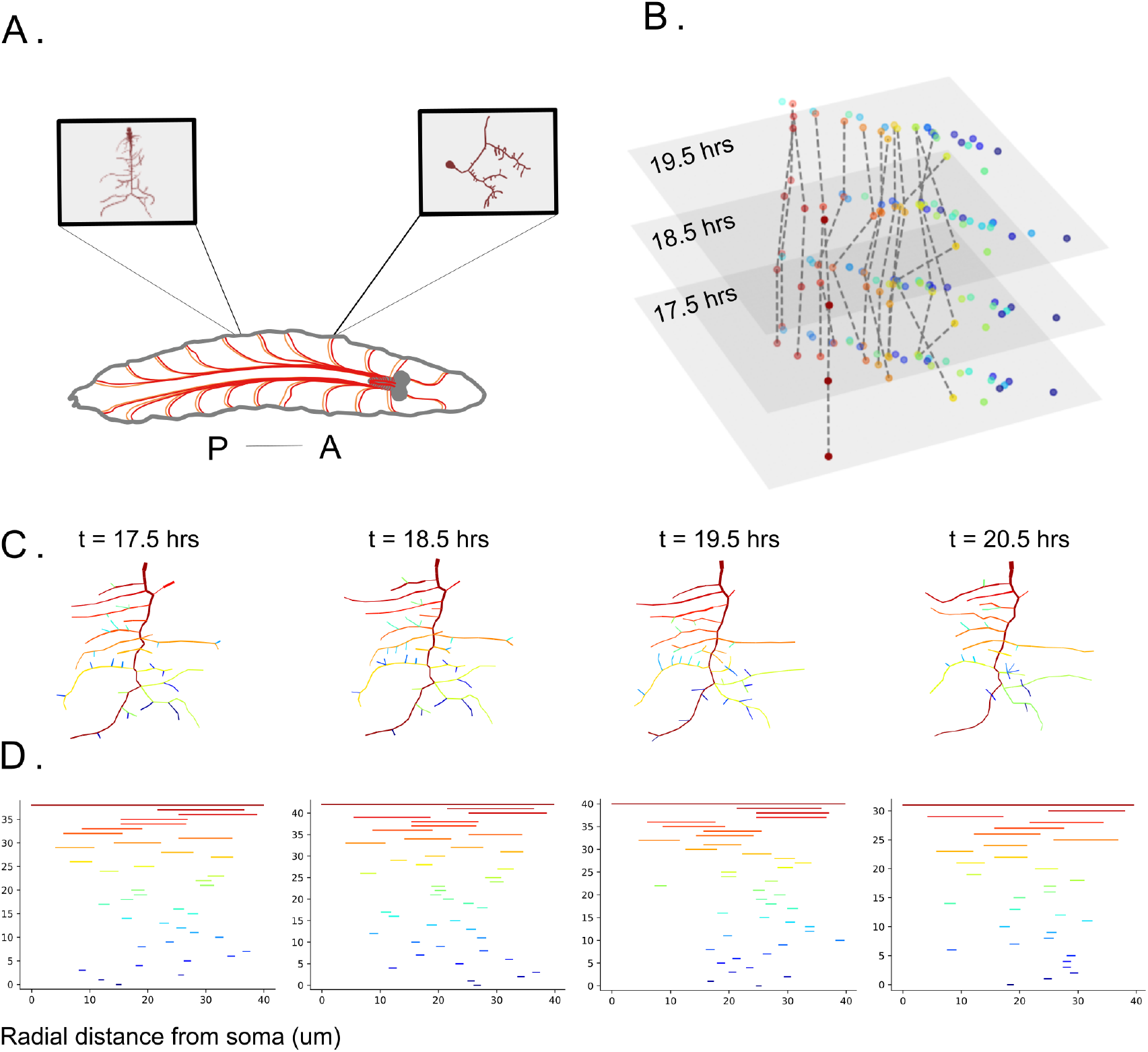
Topological characterization of neuronal development. **A**. Reconstructed morphologies of c1da (left) and c3da (right) sensory neurons in the *Drosophila* larval peripheral nervous system. P: posterior; A: anterior. **B**. Vineyard plot shows the evolution of persistence diagrams over developmental time, highlighting topological changes in neuronal structure. **C**. Example of c1da morphological reconstructions across development (from t=17.5hrs to 20.5hrs). Colors represent the branches and the bars in the persistence barcode, ranging from longer (red) to smaller branches (blue). **D**. Temporal topological morphology descriptor, represented as a series of persistence barcodes, derived from the c1da morphology reconstructions in C.

Note that due to the varying branching structure of neurons at different time steps, the persistence diagrams don’t have the same number of branches at different time points. To address this issue, while matching points between diagrams we include in the persistence diagrams the diagonal Δ = (*b*_*i*_, *d*_*i*_)|*b*_*i*_ = *d*_*i*_, as described above, which includes infinitely many points with the same birth and death times:

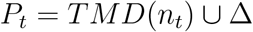

Therefore, during the matching of points between two consecutive time steps *t*_1_, *t*_2_ the points in the two diagrams can also be matched to points in the diagonal, that correspond to infinitely small branches in the tree structure.

## 3 Results

### 3.1 Dataset

Among the four classes of *Drosophila* larva sensory da neurons, Class I (c1da) proprioceptors play a crucial role in sensing body deformations, with their function closely tied to morphological adaptations [26, 32]. Their comb-liked dendrites undergo deformation during larval movement, making them particularly suitable for studying the relationship between dendritic morphology and differentiation processes [19, 58]. To analyze their developmental trajectory, we used a high-temporal-resolution dataset spanning the entire development ([19];https://doi.org/10.5281/zenodo.4290200). To visualize dendritic morphology, membrane-tagged fluorescent proteins (GFP) were expressed in c1da neurons, enabling detailed tracking of their growth dynamics [19]. This dataset covered the embryonic stage (before hatching), starting 16 hrs after egg laying (AEL), with imaging performed at the minute scale, capturing fine morphological changes throughout development. After hatching (24 hrs AEL), Class I dendrites were imaged at 30 hrs, 50 hrs, and 72 hrs AEL, providing a comprehensive view of dendritic growth as neurons expanded alongside body growth. This dataset represents one of the most complete records of the differentiation process, making it ideal for testing our method. By providing a detailed timeline of subtle morphological variations at each time point, it serves as a benchmark for evaluating the sensitivity of our approach in detecting small-scale dendritic transformations.

To further evaluate the sensitivity and robustness of our method, we leveraged the well-characterized morphology of Class III dendritic arborization (c3da) neurons in *Drosophila* larvae. C3da neurons, which respond to gentle touch and noxious cold stimuli [60], exhibit a few long primary dendrite branches decorated with numerous barbed-like, short, highly dynamic terminal branchlets that are essential for sensory function [2]. These transient and motile terminal branches introduce morphological variability, posing a challenge for topological methods, as smallscale structural changes could impact classification accuracy. We used a dataset (https://doi.org/10.5281/zenodo.6347438) that examines the distinct roles of six actin-modulatory proteins (AMPs) in shaping c3da terminal dendritic branches and the morphological effects of individual AMP mutations [54]. (1) Twinstar/Cofilin, an actin-severing protein, regulates actin dynamics at branching sites and is essential for branch formation across all da neuron classes. (2) Arp2/3, an actin-nucleating complex, transiently localizes to branching sites to initiate branchlet formation via branched actin networks. (3) Ena/VASP, a barbed-end-binding protein, promotes lateral branching in all da neuron types. (4) Singed/Fascin, an actin-bundling protein, exhibits neuron-type-specific function, being localized exclusively to the actin-rich short terminal branches of c3da neurons. (5) Spire, another actin-nucleation factor, shows differential regulation in c1da and c4da neurons. (6) Cappuccino (Capu), also an actin nucleator, regulates terminal branching specifically in c3da neurons.

This dataset includes in vivo time-lapse imaging data (GFP) with minute-scale resolution, capturing the morphology and dynamics of c3da small terminal branches in both wild-type neurons and AMP loss-of-function mutants. Using this dataset, we determine whether our method can accurately distinguish morphological variations driven by AMP mutations despite the inherent variability of highly dynamic terminal branches. This serves as a rigorous test of our method’s sensitivity to subtle, transient morphological differences, further validating its ability to classify neuronal structures based on topology.

### 3.2 Topological descriptor of growing trees

The topological description of neurons, Topological Morphology Descriptor (TMD) [35], transforms the three-dimensional shape of a neuron (Figure 1A) into a persistence barcode. The persistence barcode (Figure 1B) is a set of bars encoding the start and the end radial distances of each neuronal branch, tracing the topological lifetime of a branch. The persistence diagram (Figure 1C) is an equivalent representation in which the start and end distances are presented by a point instead of a bar. This generates a two-dimensional projection (Figure 1C) in which each point corresponds to a branch’s start and end radial distances. This representation enables the robust averaging and the vectorization of the topological representation of trees. For example, a Gaussian averaging of the points in the persistence diagram generates a persistence image [1], which can be used as input to machine learning algorithms (Figure 1D).

The Topological Morphology Descriptor (TMD) has been successfully applied to classify neurons [34] and glial cells [14], as well as for the computational generation of neuronal morphologies [36]. These applications relied on static datasets captured at a single time point—typically the mature stage—where neuronal branching patterns are relatively stable. In contrast, microglia exhibit highly variable branching, requiring population-level analysis and advanced bootstrapping techniques to achieve statistical robustness [14].

In our study, we focus on the developmental trajectory of individual neurons rather than populations, making bootstrapping techniques unsuitable. Moreover, our goal is not just to classify morphologies but to understand how dendritic structures emerge over time. Capturing temporal changes in a topological framework poses significant challenges. While multidimensional persistence homology offers one possible solution [25, 51, 38], it assumes access to continuous or dense data across dimensions. In biological imaging, however, developmental processes are only partially observed—recorded at discrete time points—resulting in fragmented datasets that are incompatible with most multidimensional persistence approaches. These constraints motivate the need for a new descriptor that integrates topological structure with temporal progression in a biologically realistic setting.

For these reasons, we extended the TMD to incorporate the dimension of time, inspired by the idea of persistence vineyards [13]. Persistence vineyards are used in topological data analysis to link the persistence diagrams of an object across an orthogonal dimension. A successful application of vineyards is the persistence homology transform [57], which represents the persistence homology of an object in different angular directions [27, 55]. We extended the classic topological description of neurons to the Temporal Topological Morphology Descriptor (TTMD). The TTMD is a collection of TMD representations of a neuronal tree at different time steps {*PD*_0_, *PD*_1_, *PD*_2_, …, *PD*_*k*_*}*. We used TTMD to analyze dendritic morphologies from the peripheral nervous system of *Drosophila* larvae at different developmental time points to quantify morphological differentiation over time (Figure 2A). For example, the *Drosophila* Class I dendritic morphologies (Figure 2C) were analyzed at discrete time steps to extract their respective persistence barcodes (Figure 2D) and persistence diagrams (PD).

The main challenge in this framework is establishing a robust correspondence between persistence diagrams across different time points. To address this challenge, we employ the Hungarian matching algorithm [39], as implemented by Munkres [46], to link each point in the persistence diagram *PD*_*t*_ at time *t* with a corresponding point in the diagram *PD*_*t*+1_ at time step *t* + 1. This matching process yields a sequence of persistence diagrams *PD*_0_ → *PD*_1_ → *PD*_2_ → · · · → *PD*_*k*_ in which links are established between the points of the diagrams at consecutive time steps (Figure 2B). The TMD-vineyards identify the most probable bars corresponding to continuously growing branches in the biological neurons, overcoming the costly process of manual tracking of the branches across time.

### 3.3 Classification of Class I and Class III neuron morphologies

To assess whether class I (Figure 3A) and class III neurons (Figure 3B) can be distinguished using topological analysis, we classified their morphologies based on their persistence images. Given the pronounced morphological differences between the two neuron types, we hypothesized that persistence images at individual time steps would effectively differentiate these classes, serving as an informative first-pass analysis. The classification results using three different methods—linear, quadratic, and tree classifiers—demonstrated that the tree classifier achieved the highest accuracy (82%), followed by the linear discriminant classifier (79%) and the quadratic classifier (58%) (Figure 3). These findings indicate that the tree classifier is the most effective approach for distinguishing between class I and class III neurons, the tree classifier was used for all subsequent classification tasks in this study.

**Figure 3:**
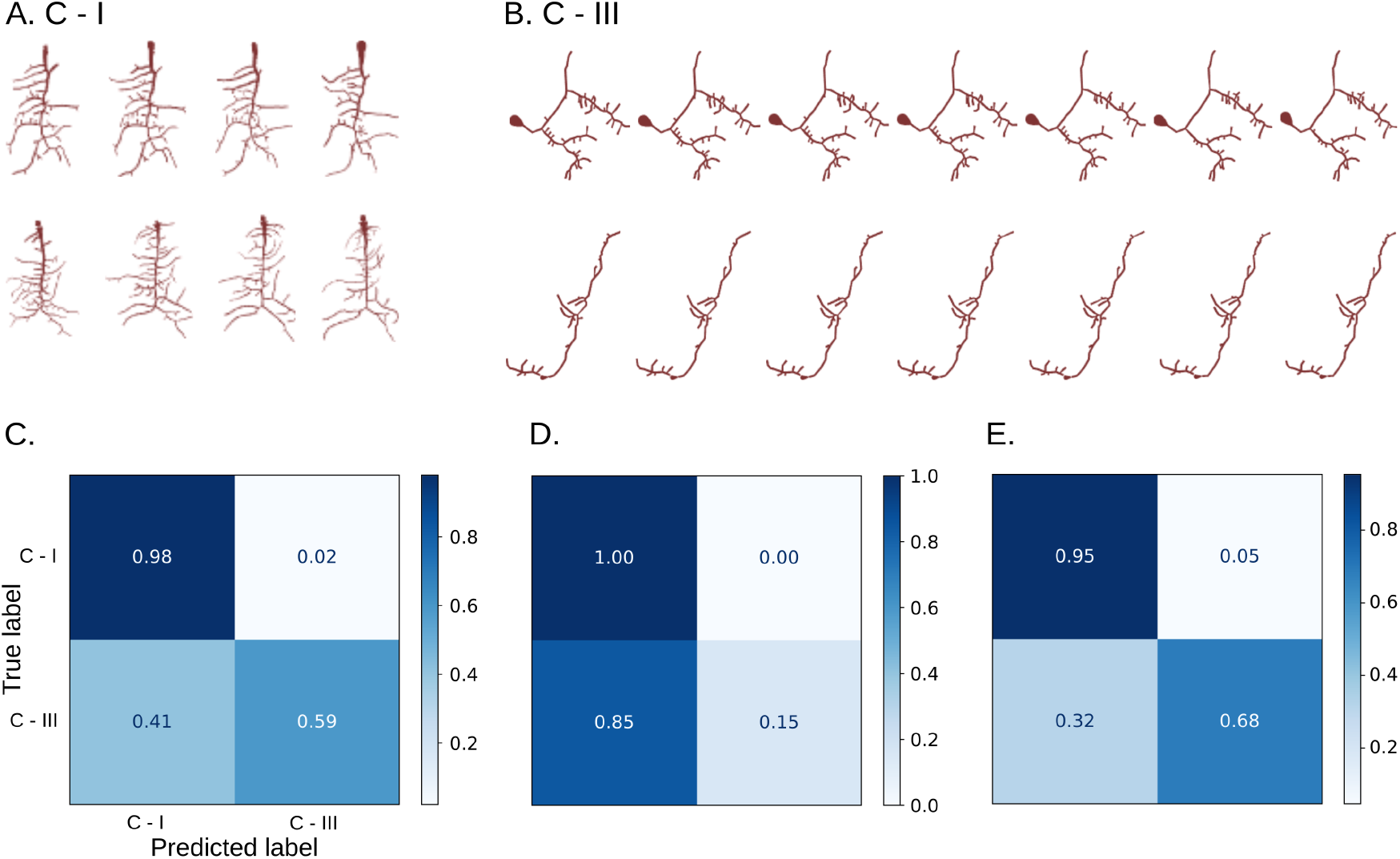
Classification of Class I and Class III neurons using topological morphology descriptors (TMDs). **A**. Representative morphological reconstructions of Class I neurons. **B**. Representative morphological reconstructions of Class III neurons. **C**. Confusion matrix for a linear classifier applied to TMD-images of the reconstructed morphologies, yielding 79% overall accuracy. Class I neurons are classified with 98% accuracy, while 41% of Class III neurons are misclassified as Class I. **D**. Quadratic classifier performance, achieving 58% overall accuracy, with high precision for Class I but poor discrimination of Class III. **E**. Tree-based classifier achieves 82% accuracy, with improved classification for Class III neurons.

**Figure 4:**
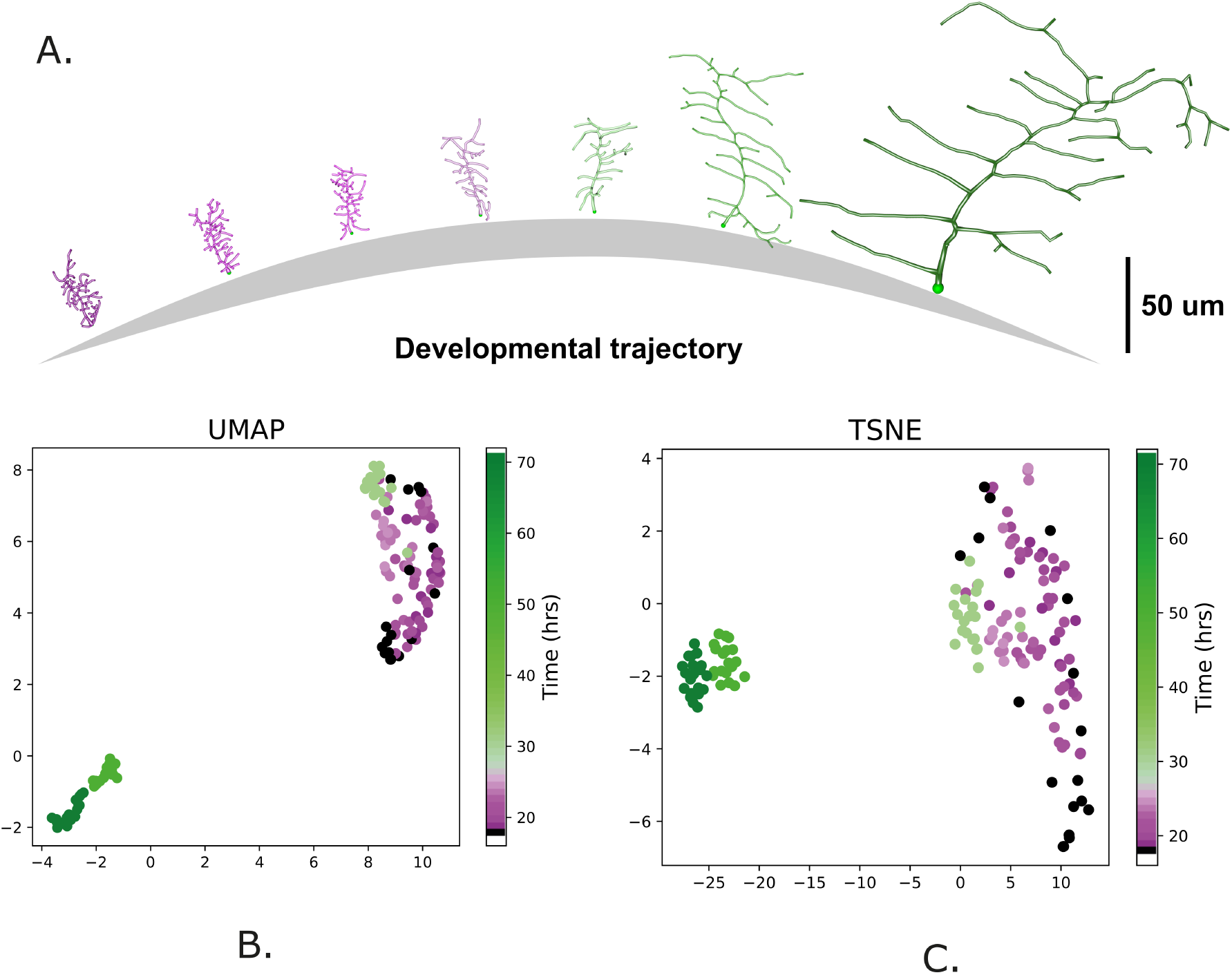
Visualization of developmental progression in Class I neurons. **A**. Representative reconstructions of Class I neurons across developmental time, illustrating morphological changes along a schematic developmental trajectory. Scale bar: 50 µm. UMAP projection of persistent morphological features (TMD) extracted from Class I neurons imaged at multiple time points. Color indicates developmental time (hours), with purple indicating embryonic stages and green representing larval stages. **C**. t-SNE visualization of the same morphological features shows even more distinct clustering corresponding to developmental stages.

Although the tree classifier demonstrated the highest accuracy among the methods tested, more advanced machine-learning techniques could further enhance performance. Approaches such as boosted gradient tree models (e.g. XGBoost) and convolutional neural networks (CNNs) are particularly well suited for capturing complex patterns and non-linear relationships in data, as shown in [37]. However, a comprehensive exploration of more complex classification methods, such as non-linear models or deep learning approaches, lies beyond the scope of the present study. Here, we deliberately focused on linear classifiers to ensure that the classification performance reflects the intrinsic sensitivity of the TMD-based representations to morphological differences, rather than potential artifacts introduced by more advanced models. Future work could leverage these advanced techniques to improve classification accuracy and potentially uncover additional insights into topological distinctions between neuronal classes.

### 3.4 TMD reveals Class I neuron developmental trajectories

Next, we focused on analyzing Class I neurons across different developmental stages. To quantitatively assess the morphological maturation of these sensory neurons, persistence images of the TMD of Class I neurons were used to generate two-dimensional embeddings through UMAP [44] and t-SNE [59], while preserving neighborhood relationships that indicate morphological similarity. Each neuron is represented as a single point in the embedding (Figure 4), with developmental time encoded using a color gradient ranging from pink (early stages) to green (late stages).

The results reveal that the topological embedding of Class I neurons aligns well with their developmental stages (Figure 4). The main developmental stages—early (pink), intermediate (light green), and late (dark green)—are more easily distinguishable than the finer variations observed within each stage. Both UMAP and TSNE plots (Figure 4) revealed that developmental time was a major factor driving variation, with neurons becoming progressively more morphologically distinct over time. During the embryonic stages (16-22 hours AEL), cells exhibited a continuous topological distribution from the early phase of primary brain development and differentiation (represented by dark and dark pink colors). In contrast, the pruning and subsequent stabilization phases were marked by lighter pink shades [19]. Post-embryonic first-instar c1da neurons (24 hours AEL) are marked in light green, positioned near embryonic neurons (Figure 3). Meanwhile, later-stage neurons (50–72 hours AEL) formed distinct clusters in darker green shades, indicating greater dendritic divergence at advanced developmental stages, as predicted by multidimensional morphometric analysis. Therefore, we demonstrate that the TMD, as a single topological descriptor, captures sufficient information to distinguish key developmental transitions in Class I neurons. Remarkably, this was achieved without the need for multiple, manually selected morphometric features, as used in previous studies to characterize growth dynamics and developmental stages of class I da neurons [19].

### 3.5 Tracking and classifying Class III neuron mutation dynamics with TTMD

To assess the sensitivity and robustness of our method, we used the well-characterized Class III dendritic arborization (c3da) neurons in *Drosophila* larvae. These neurons feature a long primary dendrite that gives rise to numerous highly dynamic branchlets, which are crucial for sensory function. The fine-scale complexity of these branchlets poses challenges for anatomical analysis of Class III neurons. The topological morphology descriptor (TMD) effectively captures this variability through the presence of small topological components near the diagonal (Figure S. 1).

Considering all available data points for each cell type, we applied a tree classifier to distinguish between the control groups and the mutant cell types. The control samples (C161, 40A, and G13) were grouped together for this analysis. Then, a classification was performed on the different mutant groups (Twinstar/Cofilin, Arp2/3, Ena/VASP, Singed/Fascin, Spire, and Capu, see also Figure S 5) and one collective control group. The high classification accuracy observed (see Figure 5A) demonstrates that the TTMD algorithm effectively differentiates the control from the mutant groups based on their morphological characteristics. Despite the increased variability introduced by genetic perturbations, the TTMD algorithm achieved a high classification accuracy between different mutants, highlighting its sensitivity in capturing structural differences associated with each mutation. These findings suggest that TTMD can reliably distinguish both subtle and pronounced morphological alterations across different phenotypes that correspond to different genetic variations.

**Figure 5:**
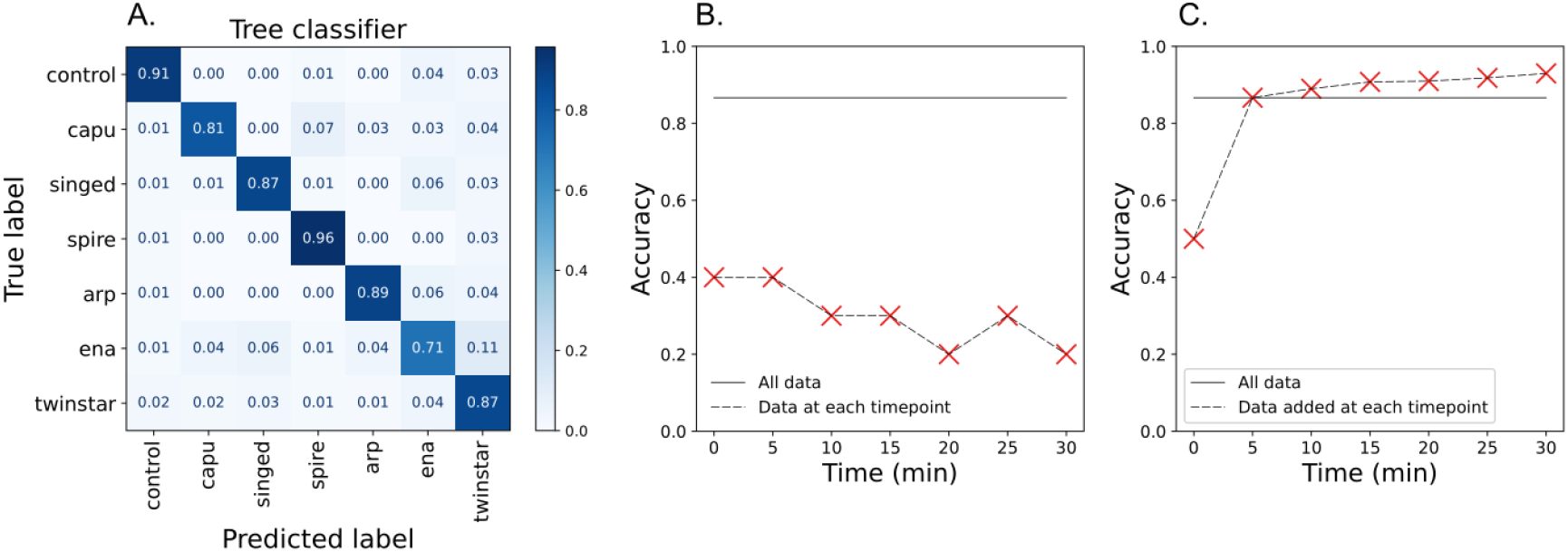
Temporal topological classification of Class III neurons. **A**. Confusion matrix showing classification results using a tree-based classifier on TTMD of Class III neurons across seven developmental time points. Overall accuracy is 86%. The control category includes three distinct control genotypes. **B**. Accuracy of independent-timepoint classification, that remains below 40%, is compared to the accuracy of full dataset classification (“All data”). **C**. Cumulative classificationmarkedly improves classification accuracy, exceeding 80% after initial timepoints. This highlights the importance of dynamic morphological descriptors for genotype discrimination.

To further investigate the developmental origins of these morphological differences, we evaluated classification accuracy across distinct time points (see Figure 5B-C). When analyzing individual time points separately using TMD (Figure 5C), classification accuracy remained consistently below 40%, indicating that the morphology at any single time step provides only partial information. In contrast, when incorporating multiple time points into the classification framework (Figure 5C), accuracy progressively increased, converging to the highest values achievable with the TTMD algorithm (see Methods Section 2.5), above 80% after only a couple of time-steps were introduced.

Although large-scale dendritic rearrangements are limited over this time scale (30 min, Figure S 5), incorporating multiple time points captures subtle, local fluctuations in dendritic geometry and branching topology that are not apparent in a single snapshot (Figure S 6). These small fluctuations, when encoded through topological descriptors, contribute additional independent features that improve the robustness of the classification (Figure 5). This analysis indicates that different mutants exhibit different temporal variability (Figure S 6), which improves separability even when the global structure remains similar (Figure S 5). Thus, the improvement in classification accuracy arises from integrating these subtle temporal signatures across time points. These results emphasize the importance of developmental trajectories in morphological differences, reinforcing the need for time-aware analytical approaches when studying neuronal morphology.

### 3.6 The developmental trajectory of a neuron through topological vineyards

In the previous classification tasks, the matching between persistence diagram across consecutive time points (Section 2.7.1) was not explicitly used. Nevertheless, we obtained high classification accuracy, indicating that precise developmental trajectories are not essential for reliably classifying neuronal types during development.

However, the matching between time steps of individual branches in the underlying neuronal structure is essential for a more comprehensive understanding of developmental processes. For instance, [19, 54] proposed specific growth mechanisms to explain the experimental observations in their datasets, by tracking the branches during development. By applying Munkres matching across time steps, we propose an automated matching that reliably approximates manually tracked branches, while substantially reducing the time required. For example, in Figure 6 we reproduced Figure 5. [19] using the Munkres matching between branches. Although TMD-vineyards do not guarantee exact anatomical identity for every matched branch, they capture the relevant population-level dynamics of branch persistence, appearance, and disappearance. Taking into account that the manual matching of branches requires a significant amount of effort and time, the statistical results achieved in a few seconds with the topological matching yield a signicant advantage in terms of methodological efficiency. The TMD-vineyard matching serves as an automated topological registration procedure. This approach facilitates the analysis of experimental datasets and supports efficient processing of the increasingly large datasets that are becoming available.

**Figure 6:**
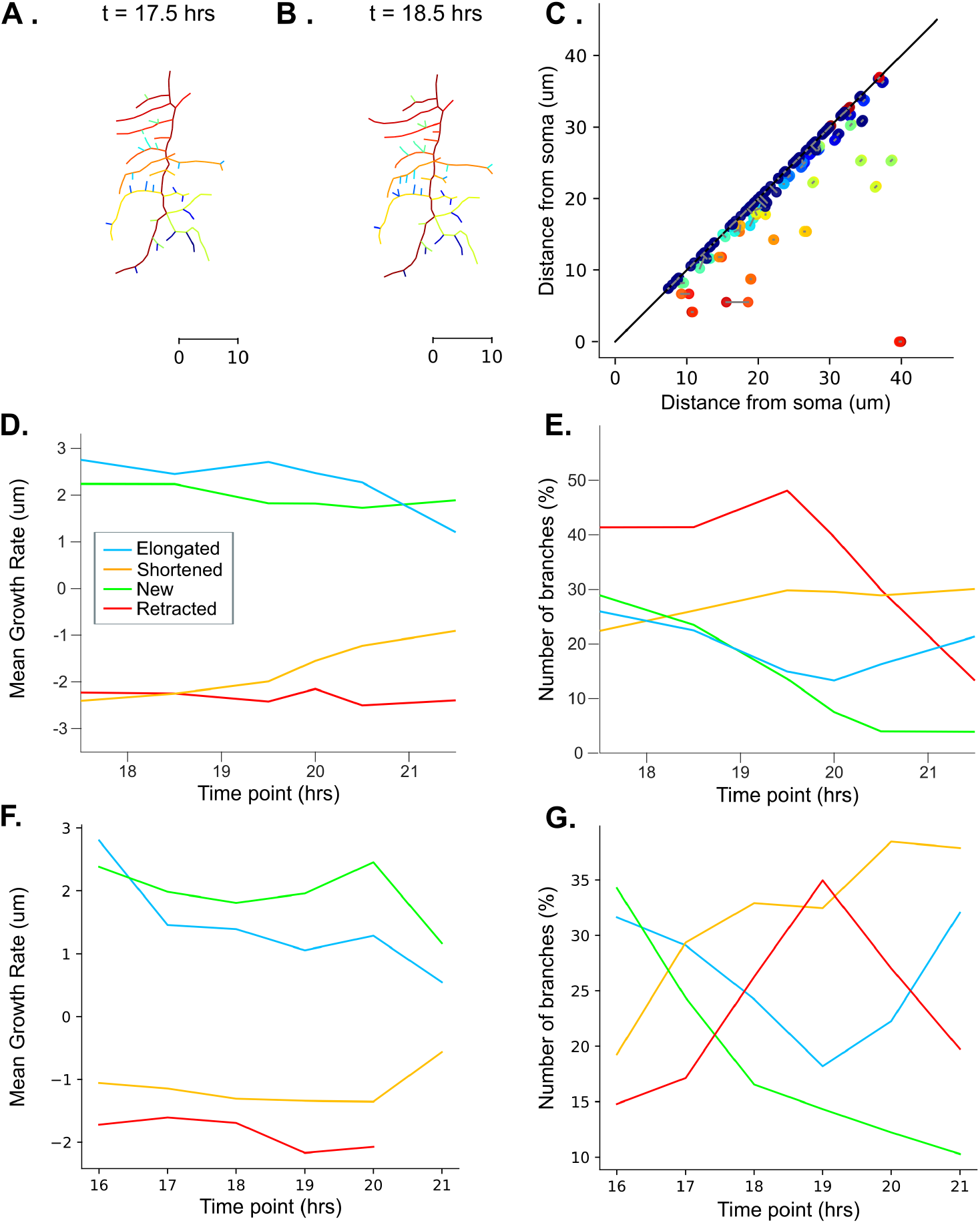
Topological branch tracking through vineyards quantifies retraction phase dynamics in Class I neurons. Example morphology with branches colored from red to blue according to their topological tracing from 17.5 hrs (**A**.) to 18.5 hrs (**B**.) of development. **C**. Topological matching of branches (A.-B.) at time steps 17.5 and 18.5. Mean branch dynamics quantified as growth rates (um / 1 hr) (**D**.) and percentage of branches (**E**.) for all branches during the different phases: elongated, shortened, new and retracted, reproduced from Fig. 5 [19]. Mean branch dynamics quantified as growth rates (um / 1 hr) (**F**.) and percentage of branches (**G**.) for all branches during the different phases: elongated, shortened, new and retracted, based on the topological matching of branches.

## 4 Discussion

In this study, we investigated the properties of branching morphologies by analyzing the differences between distinct arbor types during development and across mutants. To overcome the limitations of traditional morphometrics, such as manual feature selection biases, we extended the Topological Morphology Descriptor (TMD) to capture temporal variations in neuronal morphology [29, 31, 35]. Our results show that TMD is sufficient to identify biologically relevant growth stages in Class I dendrites, capturing key developmental transitions with a topological descriptor. However, in the case of Class III neurons, where mutations subtly alter the branching dynamics of small terminal branches while leaving the overall morphology largely intact, TMD cannot capture these subtle changes.

To address this limitation, we developed the Temporal TMD (TTMD), which incorporates temporal information, as a sequence of topological descriptors, and enables the classification of subtle morphological differences across time. TTMD successfully distinguishes mutant phenotypes in Class III neurons that would otherwise go undetected by static descriptors. Finally, we use the automated topological matching of branches across timepoints to study growth dynamics in Class I neurons. We show that using the topological matching of branches, which substantially improves computational efficiency relative to manual branch matching, we capture the relevant population-level dynamics of branch persistence, appearance, and disappearance.

By providing a computationally efficient and scalable alternative to multidimensional morphometric approaches, our method enables a more robust analysis of neuronal development and arborization patterns [35, 8]. This advancement not only enhances our understanding of dendritic growth and neural circuitry formation but also lays the foundation for future research into developmental mechanisms and neural disorders, particularly with the increasing availability of high-throughput and high-temporal-resolution imaging techniques.

### 4.1 Limitations of the study

While our method provides a powerful framework for analyzing neuronal morphology, we acknowledge a few limitations. As imaging techniques advance, allowing for higher temporal resolution and the capture of faster dynamics—such as those occurring on the second time scale—morphological differences and branch morphogenesis will become increasingly nuanced. At such fine temporal scales, distinguishing biologically relevant structural changes from measurement noise and imaging artifacts may pose a challenge. Additionally, smaller and more transient morphological features may introduce variability that could affect the robustness of our approach. Future refinements should focus on enhancing sensitivity to these rapid structural dynamics and improving noise handling in high-temporal-resolution datasets. Nevertheless, even at the minute-scale resolution of our data, our method successfully differentiated Class III neuron mutants that specifically affected the highly motile small terminal branches, demonstrating its strong sensitivity to subtle morphological changes and its potential for application in even more temporally refined datasets.

One assumption of our method is that dendritic topology alone provides sufficient information for classifying morphologies. While topology captures the branching architecture and overall structural organization, it does not account for differences that are orthogonal to topology, such as dendritic thickness or subtle geometrical variations in cytoskeletal structure. If two neuronal populations share the same topological structure but differ in properties like branch diameter, our method will not effectively distinguish them. Future work could integrate complementary morphological descriptors, such as volumetric or thickness features, to enhance classification capabilities. Despite this limitation, our approach remains highly effective in distinguishing morphologies where branching patterns play a key role, offering a scalable and computationally efficient alternative to more complex multidimensional morphometric analyses.

### 4.2 TTMD and existing morphometrics

Our findings align with previous research, demonstrating that our method can accurately retrieve developmental stages while relying on a single topological measure rather than multiple morphometric descriptors [19, 6]. Compared to traditional approaches, the TTMD approach offers greater efficiency robustness, and flexibility across datasets. Although prior studies often required a large number of morphometric computations to uncover subtle differences, our approach even effectively distinguished mutants in Class III neurons using only one measure, highlighting the sensitivity of topological features over geometric descriptors [54]. Moreover, by providing a holistic and unbiased view of developmental stages and mutant differentiation, our method facilitates a more thorough investigation of dendritic growth patterns and branching dynamics [19, 6, 54]. Given that neuroanatomists frequently rely on the manual selection of morphometrics, which can introduce biases and overlook key structural features, our approach presents a scalable and automated alternative.

In general, the TTMD approach provides a fast, robust, and scalable solution for high-throughput time-lapse analyses of neuronal morphology, addressing the growing demand for efficient and sensitive measurement methods as imaging datasets grow larger [29, 3, 45]. By integrating our method into an open-access Python library, we ensure broad accessibility for researchers across various disciplines. Additionally, our approach holds promise for electron microscopy (EM) developmental datasets, where large acquisition gaps pose challenges for temporal alignment. By leveraging topological analysis, our method provides a cost-effective means of bridging these gaps, enabling better classification of cell types across developmental time points [20]. Furthermore, the versatility of our method extends beyond neuroscience, offering a generalizable framework for analyzing the dynamics of branching structures, such as river systems, road networks, and other biological organs such as the mammary glands and kidneys [24].

### 4.3 Conclusion

In conclusion, we present an interpretable framework for quantifying subtle differences in neuronal morphology, highlighting the importance of dendritic topology in distinguishing neuronal cell types and developmental states. Our approach is particularly well suited for developmental datasets, where structural differences are often difficult to capture using conventional morphological metrics.

An important challenge moving forward is to determine whether the observed topological differences reflect biologically meaningful heterogeneity and how they relate to neuronal function, circuit formation, and behavior. To address this, we aim to integrate automated topological branch matching with quantitative models of branch birth–death dynamics [36], building a framework similar to the time-lapse imaging of axonal arborization in zebrafish [11]. In particular, we will investigate how local branch dynamics, including formation, stabilization, and retraction [19], give rise to the global topological patterns identified here. This framework has the potential to accelerate the analysis of large developmental datasets and provide mechanistic insights into neuronal morphogenesis. Moreover, insights gained from these analyses can shed light on the role of morphologies in pathological conditions, contributing to a more comprehensive understanding of neurodevelopmental and neurodegenerative disorders.

## 6 Acknowledgments

We thank Professor Kathryn Hess for valuable early discussions that contributed to the development of this project.

