## Supplementary material for "Temporal topology provides an interpretable framework for neuronal morphogenesis"

### Supplementary Information

| Neuron class | Type | Mutant | Number of time series | Number of dendrites per series |
|---|---|---|---|---|
| Class I | Embryonic | No | 25 | 2 to 7 |
| | Larval | No | 20 | 3 |
| Class III | Control | No | 7 | 10 |
| | Capu | Yes | 7 | 10 |
| | Singed | Yes | 7 | 10 |
| | Spire | Yes | 7 | 10 |
| | Twinstar | Yes | 7 | 10 |
| | Ena | Yes | 7 | 10 |
| | Arpc | Yes | 7 | 10 |

Table S 1: Summary of the nature of all considered neuron classes, as well as the data considered for each of them. Class I neuron data come from [19] and class III ones from [54]. The embryonic datasets *t5_30*, *t6_30* as well as the six *tl_nov_friday* were excluded during the preprocessing phase.

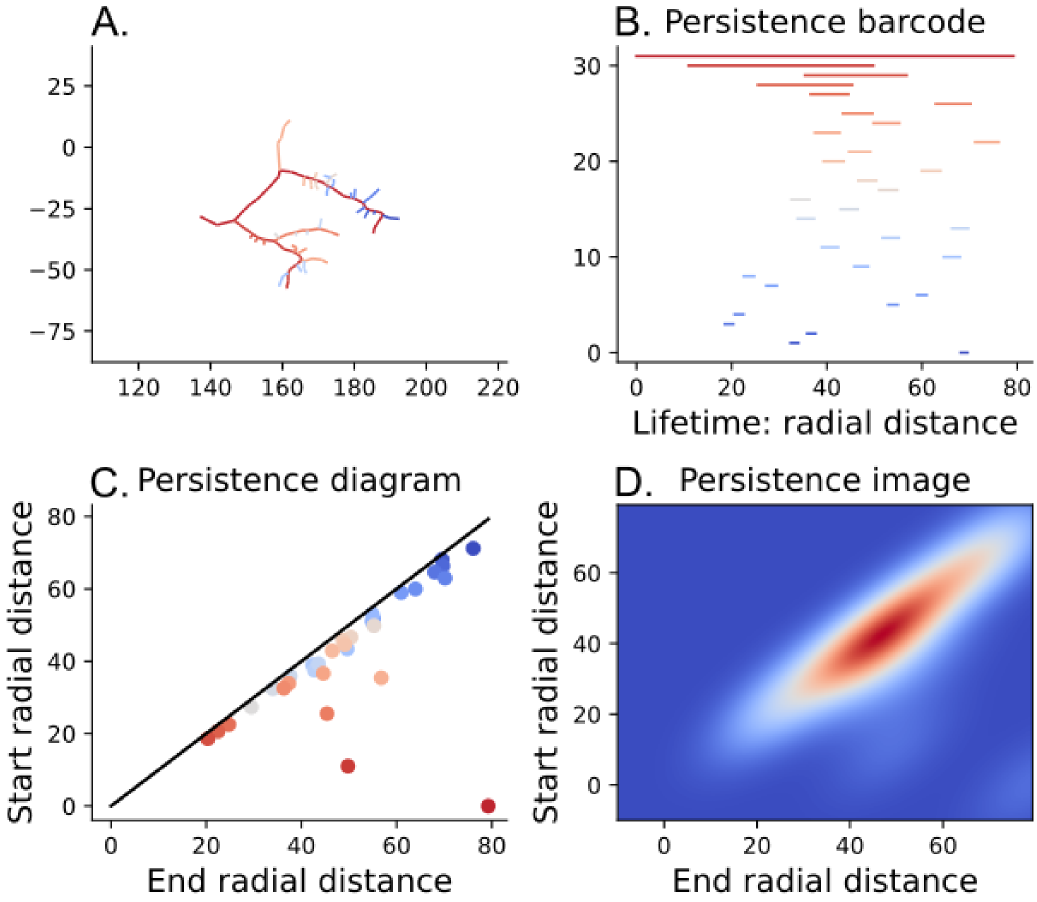


Figure S 1: **Topological Morphology Descriptor of a Class III neuron. A.** The reconstructed dendritic tree. Colors represent the branches and the bars in the persistence barcode, ranging from longer (red) to smaller branches (blue). **B.** The TMD barcode is produced through filtration of the apical graph using radial distances (in micrometers). **C.** The persistence diagram presents the same information plotted in two-dimensional space, where axes represent the birth and the death of each segment of the bar code. **D.** The persistence image shows a Gaussian kernel density estimate applied to the persistence diagram in C.

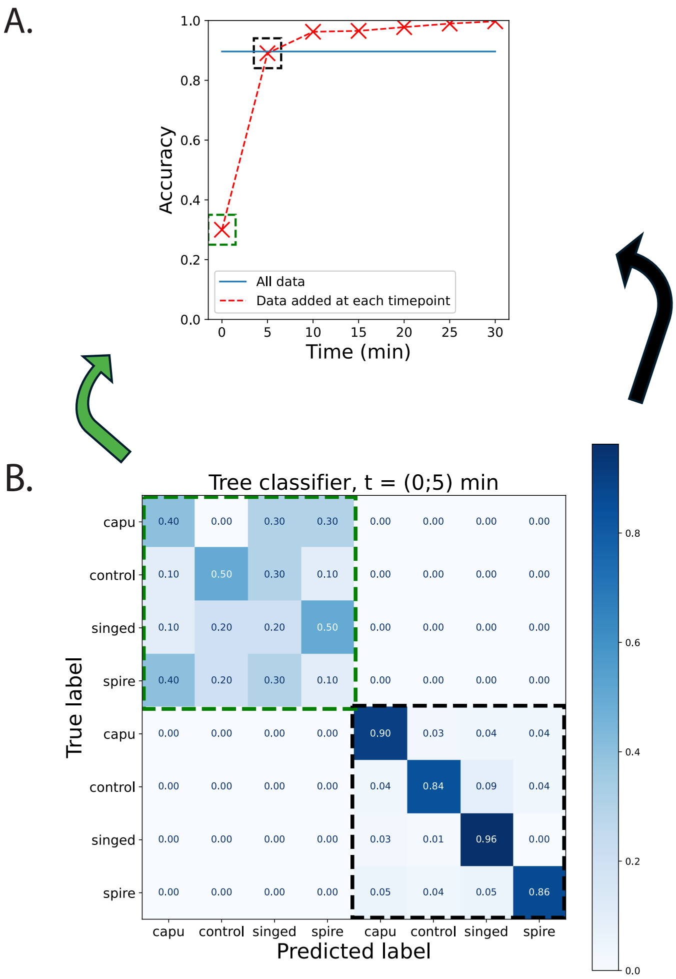


Figure S 2: **Visualization of classification dynamics for mutant Class III neurons using TTMD. A.** Classification accuracy over time with temporal integration ("memory"), showing how performance improves as data from successive timepoints are added. Red crosses indicate accuracy at each timepoint when added incrementally, while the blue line represents classification using the full dataset across all timepoints. **B.** Confusion matrix corresponding to classification at early timepoints (0–5 minutes), using a tree classifier. The matrix is divided into two main blocks: early classification performance (green dashed box) and overall performance when additional timepoints are incorporated (black dashed box). Each diagonal block corresponds to accuracy at a specific timepoint, directly linked to the accuracy curve in A.

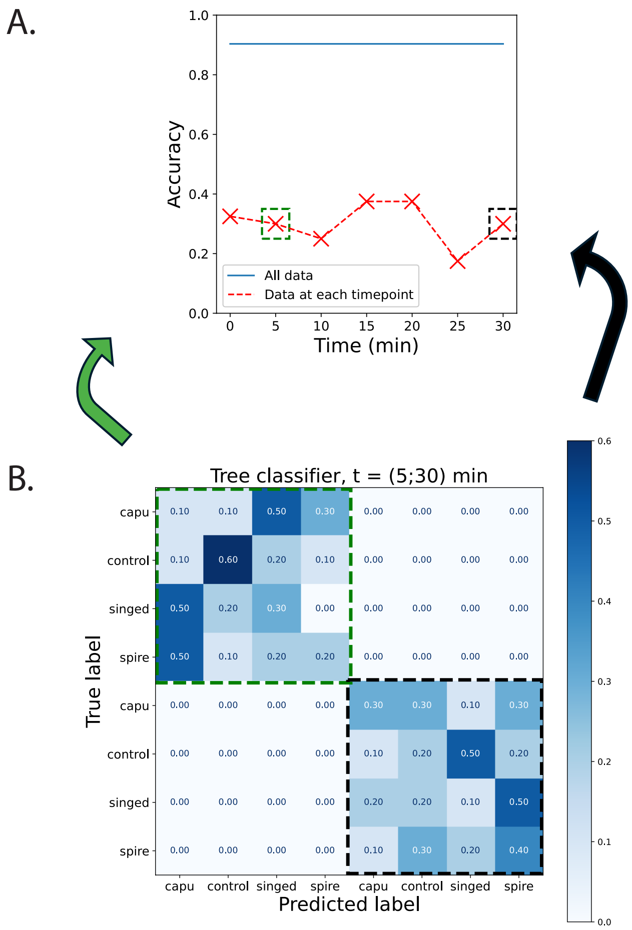


Figure S 3: **Classification performance for mutant Class III neurons without temporal memory. A.** Accuracy over time when classifiers are trained independently at each timepoint without incorporating temporal context. Red crosses show per-timepoint accuracy, and the black dashed squares highlight specific timepoints linked to confusion matrix blocks in B. The blue line shows accuracy using all timepoints combined. **B.** Confusion matrix from classification using data from two timepoints (5 and 30 minutes), trained with a tree-based classifier. The matrix is divided into two diagonal blocks: the upper-left (green dashed box) represents classification using only early timepoints, while the lower-right (black dashed box) reflects classification using later timepoints. Each block demonstrates limited performance when temporal continuity is excluded.

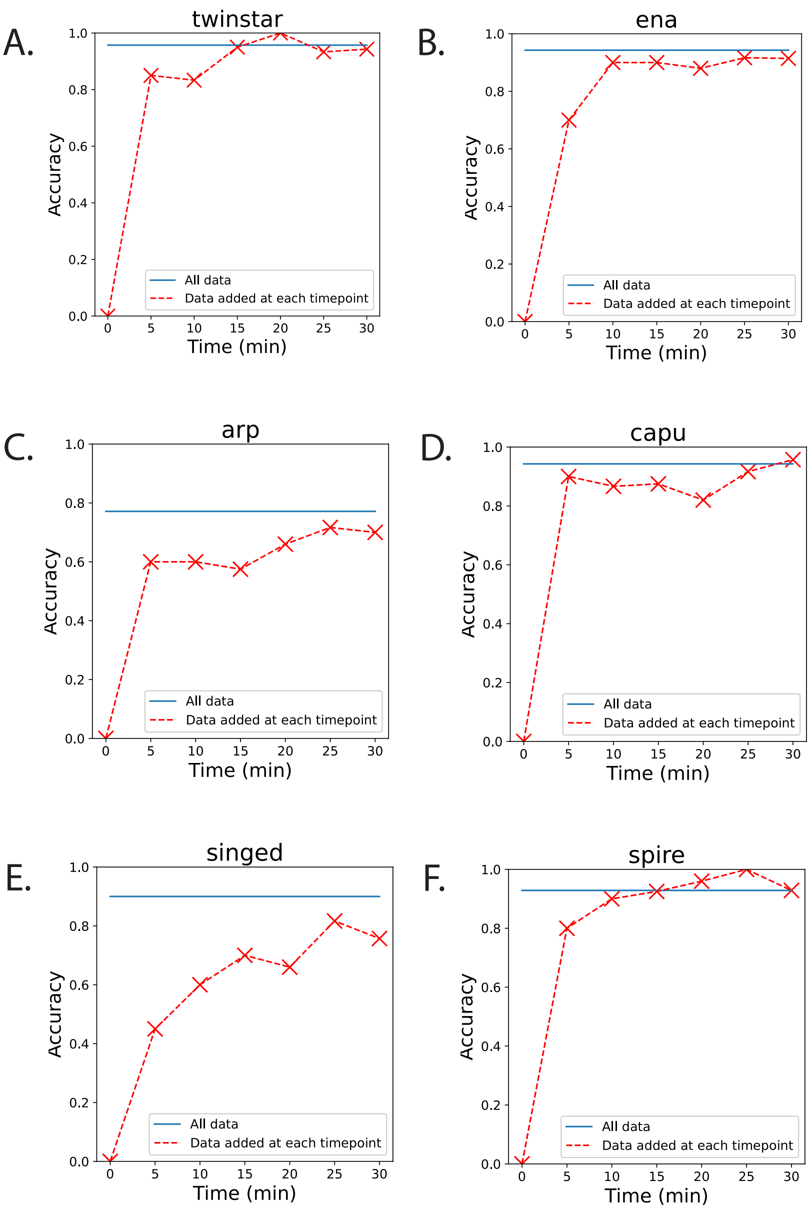


Figure S 4: **Classification accuracy across timepoints for individual Class III mutant genotypes.** Each panel (A–F) shows the classification accuracy of TTMD-based classifiers for a specific mutant line as temporal data accumulates. Red dashed lines with crosses represent accuracy when data is incrementally added at each timepoint; the blue line indicates accuracy when using the full dataset across all timepoints. **A.** *twinstar*, **B.** *ena*, **C.** *arp*, **D.** *capu*, **E.** *singed*, **F.** *spire*. Most mutants show a steep improvement in classification accuracy within the first 10 minutes, highlighting the effectiveness of temporal information in genotype discrimination.

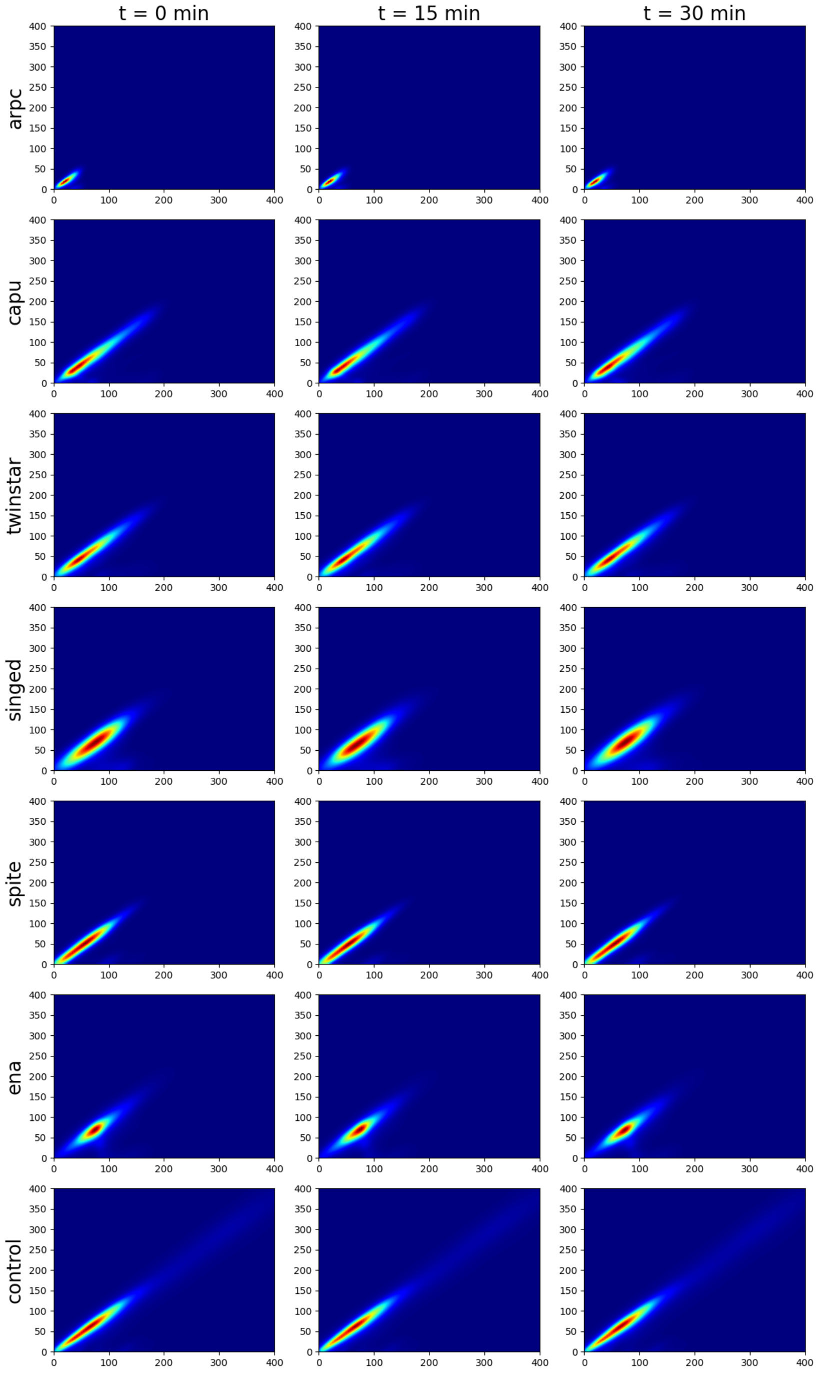


Figure S 5: **Persistence images across timepoints for Class III mutant genotypes.** Persistence images averaged over the cells of mutant types: *arpc*, *capu*, *twinstar*, *singed*, *spire*, *ena*, and *control* for three different timepoints ($t = 0$ min, $t = 15$ min, $t = 30$ min).

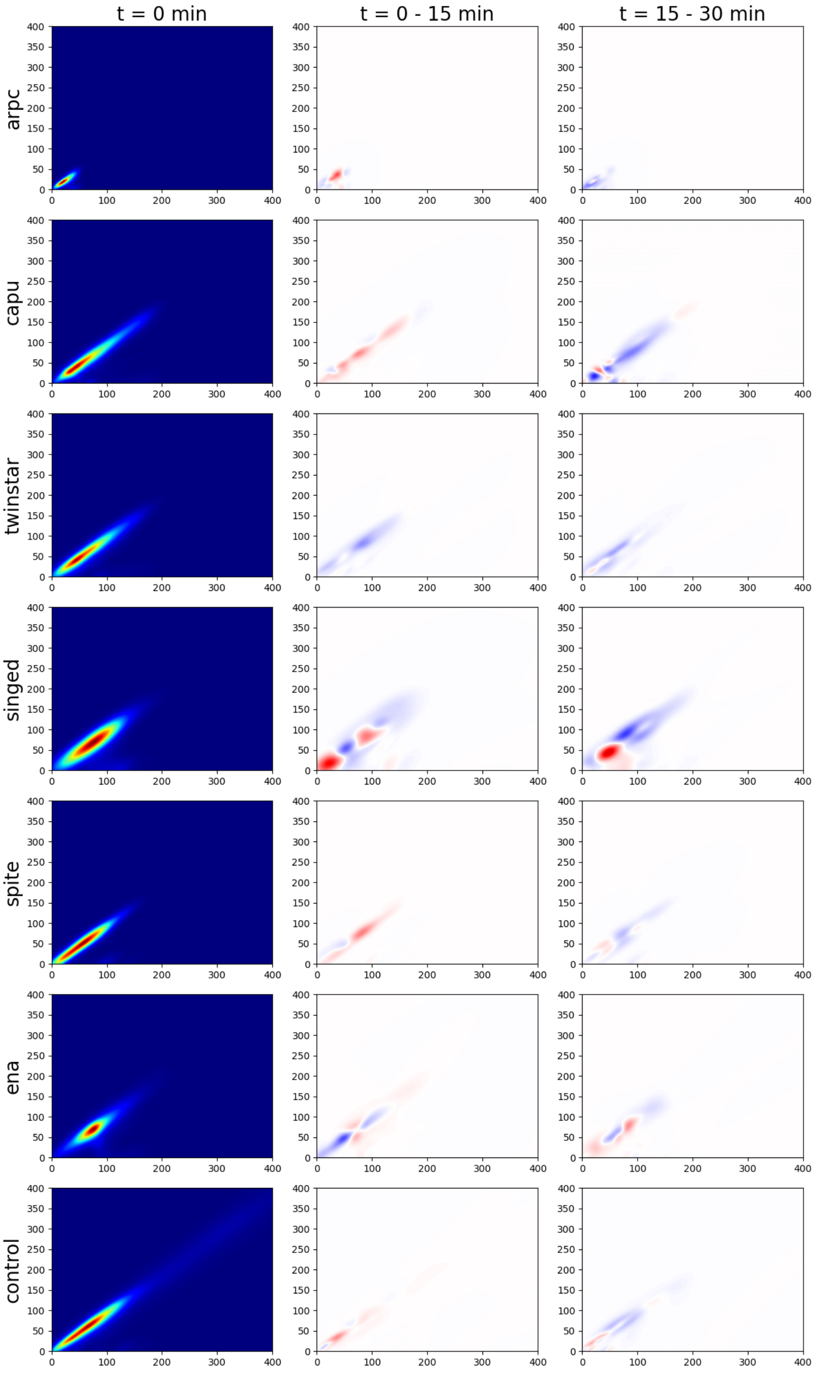


Figure S 6: **Persistence images and differences for Class III mutant genotypes across timepoints.** Persistence images averaged over the cells of mutant types: *arpc*, *capu*, *twinstar*, *singed*, *spire*, *ena*, and *control* for timepoint $t = 0$ min, and persistence image differences between timepoints $t = 0 - 15$ min and $t = 15 - 30$ min.
